# Impact of upstream landslide on perialpine lake ecosystem: an assessment using multi-temporal satellite data

**DOI:** 10.1101/808824

**Authors:** Paolo Villa, Mariano Bresciani, Rossano Bolpagni, Federica Braga, Dario Bellingeri, Claudia Giardino

## Abstract

Monitoring freshwater and wetland systems and their response to stressors of natural or anthropogenic origin is critical for ecosystem conservation.

A multi-temporal set of 87 images acquired by Sentinel-2 satellites over three years (2016-2018) provided quantitative information for assessing the temporal evolution of key ecosystem variables in the perialpine Lake Mezzola (northern Italy), which was suffered from the impacts of a massive landslide that took place upstream of the lake basin in summer 2017.

Sentinel-2 derived products revealed an increase in lake turbidity triggered by the landslide that amounted to twice the average values scored in the years preceding and following the event. Hotspots of turbidity within the lake were in particular highlighted. Moreover both submerged and riparian vegetation showed harmful impacts due to sediment deposition. A partial loss of submerged macrophyte cover was found, with delayed growth and a possible community shift in favor of species adapted to inorganic substrates. Satellite-derived seasonal dynamics showed that exceptional sediment load can overwrite climatic factors in controlling phenology of riparian reed beds, resulting in two consecutive years with shorter than normal growing season, and roughly 20% drop in productivity according to spectral proxies: compared to 2016, senescence came earlier by around 20 days on average in 2017 season, and green-up was delayed by up to 50 days (20 days, on average) in 2018, following the landslide.

The approach presented could be easily implemented for continuous monitoring of similar ecosystems subject to external pressures with periods of high sediment loads.

## 1. Introduction

Assessing the status of freshwater and wetland ecosystems and its temporal evolution in response to external events of natural or anthropogenic origin occurring in the watershed is a key requirement towards ecosystem conservation. In particular, monitoring aquatic plant communities in these systems is crucial because of their role in biogeochemical processes in water column and sediments (Schindler and Scheuerell, 2002; Jeppesen et al., 2012), their interactions with other autotrophic and heterotrophic organisms (Timms and Moss, 1984; van Donk and van de Bund, 2002; Bolpagni et al., 2014), and the provision of food and shelter to animal communities, such as fish and birds (Johnson and Montalbano, 1984; Carpenter and Lodge, 1986; Wang et al., 2017).

Satellite data can provide valuable information on a range of ecosystem processes (Hestir et al., 2015; Zhang et al., 2017; Murray et al., 2018). For more than five decades, Earth Observation (EO) techniques have been used to study inland water quality and conditions (see Tyler et al., 2016), such as: changes in turbidity (Olmanson et al., 2008), lake hydrodynamics (Pinardi et al., 2015), tidal effects on suspended solids (Eleveld et al., 2014), bloom dynamics (Duan et al., 2014), submerged macrophytes cover (Bresciani et al., 2012). EO techniques have been also used to investigate ecosystem degradation drivers, especially focusing on vegetation status. Even though literature on this topic has principally dealt with terrestrial biomes (Smith et al., 2014, and references therein), few works exist that, highlighting limitations due to satellite data features (i.e. spatial, temporal, spectral resolutions, and data availability), have addressed applications in inland and transitional environments, such as: loss and recovery of coastal vegetation after a tsunami (Villa et al., 2012), disturbance on seagrass populations in Australia (Lyons et al., 2013; Kilminster et al., 2015), turbidity-driven degradation of coral reef habitats (Fabricius et al., 2014), or aquatic vegetation changes with increased eutrophication in China (Zhang et al., 2016).

New generation EO platforms, such as Landsat 8 and Sentinel-2, host mid-resolution (10-30 m pixel side) optical multi-spectral sensors fine enough to monitor environmental phenomena at spatial and temporal scales (see Pahlevan et al., 2017) not possible before they became operational, in 2013. Such technical capabilities can improve existing EO based products and enable a number of new applications for remote sensing of aquatic ecosystems (Murray et al., 2018), some of which were recently explored, focusing either on aquatic vegetation (Stratoulias et al., 2015; Gao et al., 2017; Villa et al., 2018; Ghirardi et al., 2019), or on water quality (Dörnhöfer et al., 2016; Pinardi et al., 2018; Fritz et al., 2019; Sòria-Perpinyà et al., 2020).

The aim of this work is to expand the range of EO applications in monitoring inland water systems by providing a comprehensive framework that joins water quality and aquatic vegetation in a unique, real case scenario, i.e. using Sentinel-2 data to study the effects of an upstream landslide event that altered sediment inflow and deposition in a perialpine lake ecosystem (Lake Mezzola, northern Italy). Few works have been carried out in the last decade on the impact of altered sediment deposition on perialpine lakes dynamics, e.g. in response to upstream hydropower operations (Finger et al., 2006; 2007), or as reaction to increased flood frequency (Fink et al., 2016), and EO data and techniques potentially satisfy the requirements for becoming key tools towards further knowledge advancement on this topic.

The objective of this work is two-fold: i) to demonstrate the feasibility of monitoring the temporal evolution of key ecosystem variables in a perialpine lake at fine spatial scale (10-20 m), using Sentinel-2 satellite data; and ii) to use Sentinel-2 derived maps to quantitatively assess the impacts of an upstream landslide aftermath on the aquatic and wetland plant communities in Lake Mezzola.

## 2. Study area

Lake Mezzola (46°13’ N, 9°26’ E, 199 m a.s.l.) is a deep (average depth: 26 m, maximum depth: 69 m) perialpine lake located in northern Italy (Fig. 1). The lake, considered monomictic, covers an area of 5.85 km^2^ and has the Mera River as main tributary, with its source in the Swiss Alps (2849 m a.s.l.), 40 km North-East from the lake. The Mera River inflow, carrying sediments from the upstream Alpine basin, is responsible for seasonal periods of high turbidity and suspended solids contribution (Secchi disk depth varying between 1 and 5 m), while phytoplankton biomass is generally low (Chl-a concentration in the range 3-7 µg l^−1^), with diatoms as dominant group (ARPA Lombardia, 2019). Lake Mezzola is connected southwards to Lake Como by the Mera River, which in between the two forms a small shallow water basin, called Lake Dascio (Fig. 1).

**Figure 1.**
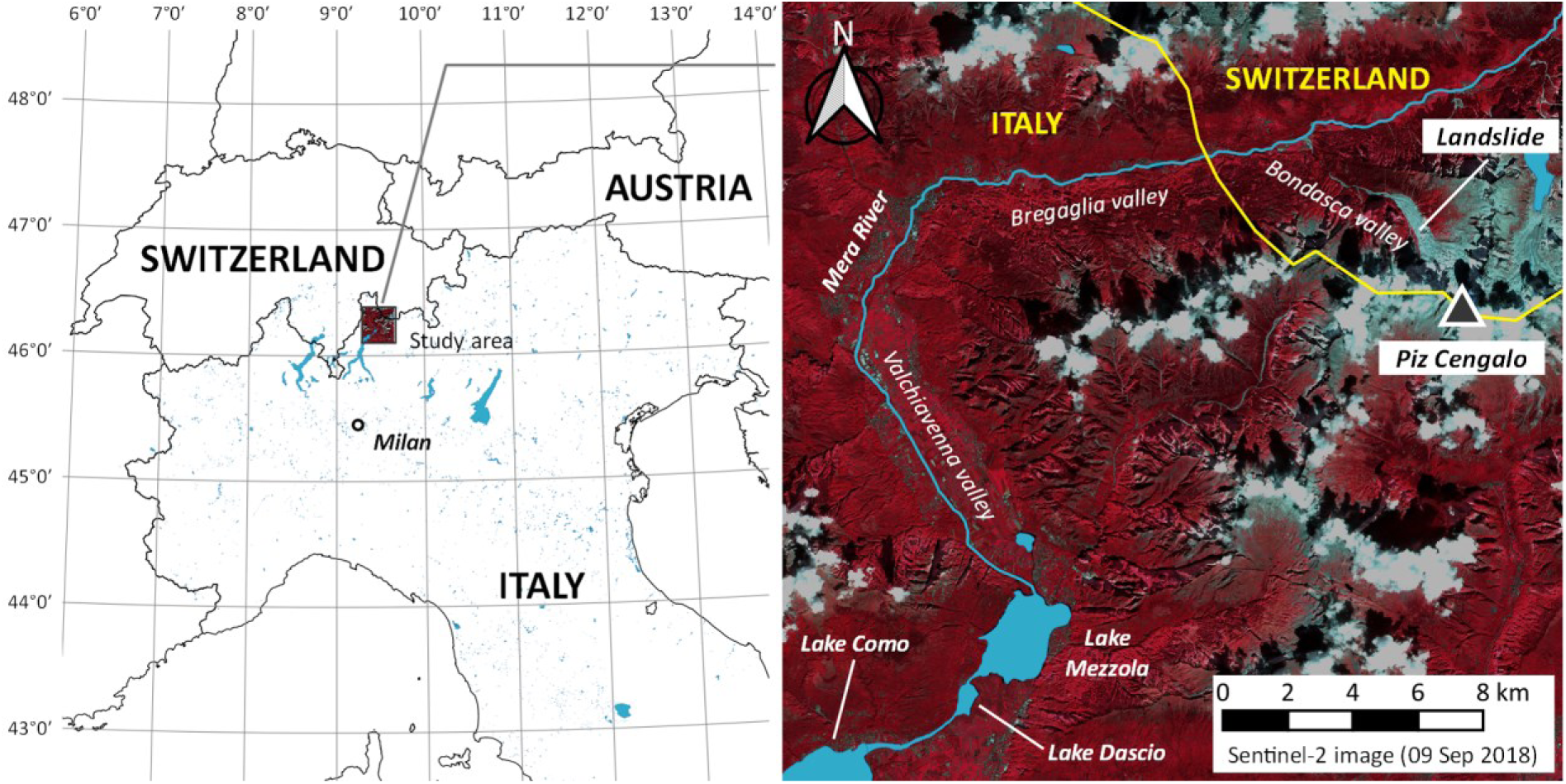
Study area overview, including a near infrared colour composite of the area derived from the Sentinel-2 image acquired on 09 September 2018.

The nature reserve of “Pian di Spagna-Lago di Mezzola” encompasses the lake and around 16 km^2^ of alluvial floodplain, called Pian di Spagna, that lies between Lake Mezzola and the larger Lake Como. The nature reserve, one of the oldest Ramsar sites in Italy and part of the European ecological network “Natura 2000”, is an important migration and breeding area for more than 200 species of birds, including a variety of waterbirds and raptors, and provides spawning and nursery areas for the rich fish fauna of Lake Mezzola.

This wetland system is characterized by lentic waters and marshes. In the southern portion of Lake Mezzola, aquatic vegetation is mainly constituted by submerged vascular macrophytes, *Groenlandia densa* (L.) Fourr., *Potamogeton perfoliatus* L. and *P. lucens* L., and benthic macroalgae *Chara virgata* Kützing and *C. globularis* Thuiller. The helophyte communities of Pian di Spagna wetland are dominated by common reed (*Phragmites australis* (Cav.) Trin. Ex Steud), with some sparse presence of *Schoenoplectus lacustris* (L.) Palla. More or less dense patches of isoetids - *Littorella uniflora* (L.) Asch, *Eleocharis acicularis* (L.) Roem. & Schult, *Limosella aquatica* L., *Elatine hydropiper* L. and *Ranunculus reptans* L. - inhabit the lakeshore area where seasonal water level fluctuations periodically leave the substrate uncovered, especially at the end of summer. In the surroundings, wet meadows and sedges can be found together with some pastures and stretches of open woodland (*Salix* spp., *Alnus glutinosa* (L.) Gaertn.).

### 2.1 Landslide event

On 23 August 2017, a major rock landslide detached from the north face of Piz Cengalo towards Bondasca valley, in South East Switzerland (Fig. 1), with a volume estimated at 3.1 million m^3^. After hitting a glacier, the event evolved into a rock and ice avalanche, finally turning into a debris flow that reached the Bregaglia valley in minutes (Mergili et al., 2019). The landslide was followed by a minor intensity event on 25 August. Because of the joint events, 150 people were displaced from the village of Bondo (Switzerland), which was partially destroyed. Resulting detrital material firstly reached the dammed river reservoir of Villa di Chiavenna (Italy), which was emptied in July as a prevention measure. Afterwards, flowing downstream in the Mera River, finer particles of landslide debris reached Lake Mezzola, triggering a massive increment in water turbidity and suspended solids loads that lasted for some months.

Following the events, the Regional Environmental Protection Agency of Lombardy (ARPA Lombardia) has put in place a specific monitoring program of the Mera River watershed and Lake Mezzola, in order to assess the environmental effects of landslide aftermath, and in particular the impacts on water quality in Lake Mezzola.

## 3. Dataset

### 3.1 Satellite data

Time series of Sentinel-2 data acquired over the study area (tile: T32TNS) covering three full years (2016-2018) were gathered. The mission is managed by the European Space Agency (ESA) and the EU Copernicus programme and data are distributed under a free and open data policy. Sentinel-2 is a constellation of two satellites: Sentinel-2A (S-2A, launched on 23 June 2015) and Sentinel-2B (S-2B, launched on 7 March 2017). Both satellites carry on board the MultiSpectral Instrument (MSI), a mid-resolution (10-60 m) multi-spectral (13 bands) push-broom imaging sensor (Drusch et al., 2012). Revisit time for Sentinel-2 is 10 days with S-2A only (up to July 2017), and 5 days with both S-2A and S-2B satellites (i.e. from July 2017).

All the available S-2A and S-2B scenes acquired from 01 January 2016 to 31 December 2018, with cloud cover less 50% on the area of Lake Mezzola and surroundings, were collected. The resulting time series (Fig. 2) was composed of 87 dates, acquired during the three years: 18 dates in 2016, 30 dates in 2017, and 39 dates in 2018.

**Figure 2.**
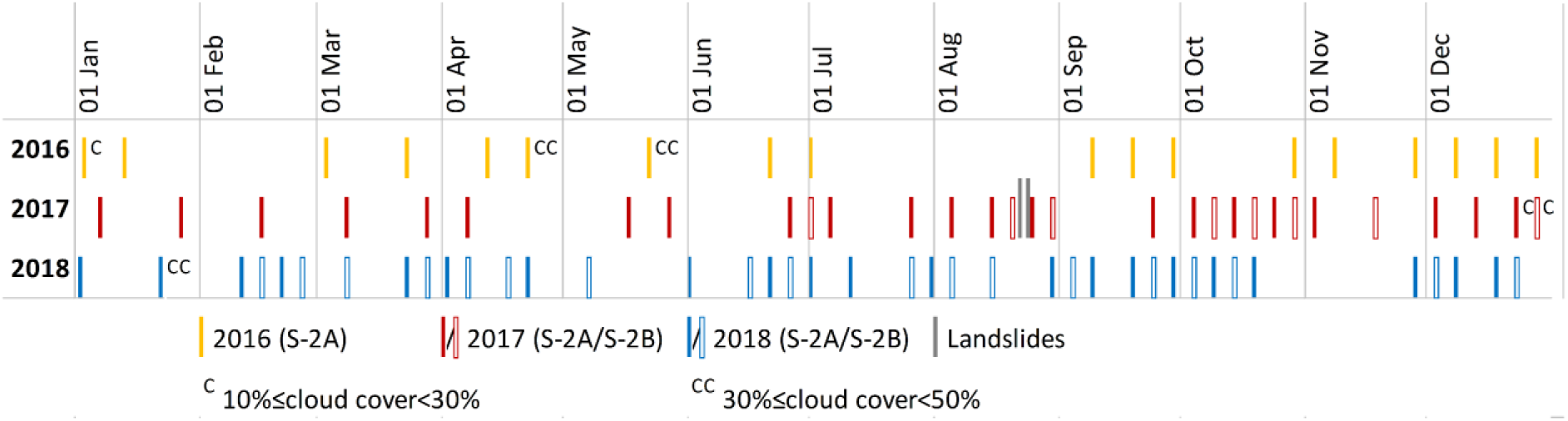
Satellite dataset, highlighting dates with cloud cover over more than 10% of the study area.

### 3.2 Fieldworks and supporting data

Time series of principal hydrological and meteorological parameters available for the last two decades were retrieved from data collected by automatic sampling stations located nearby the study area and managed by ARPA Lombardia. Daily average water level data of the Mera River collected in Lake Como (46°10’11” N, 9°23’37” E), 3 km downstream of Mera River outlet and available since 2013, were considered as good proxy of Lake Mezzola water level variation. Meteorological daily data (1994-2018) were collected from a weather station located 4 km upstream of the lake (46°14’01” N, 9°25’36” E), comprising: minimum, average and maximum air temperature, in (°C), average and maximum global radiation (W m^-2^), and cumulated rainfall (mm).

An *in situ* floristic survey was performed at the beginning of August 2018, exploring in particular the common reed and isoetid vegetation along the shoreline of the southern sector of Lake Mezzola. Afterwards, a boat-based sampling campaign was carried out on 26 September 2018, for assessing the conditions of submerged macrophytes and riparian reeds in the same area. Georeferenced photos (Camera Olympus Stylus TG-4 Tough) and direct observations of vegetation conditions were collected in order to document plant species present, reed density and phenological stage, as well as possible traces of damage.

The area covered by reed beds along the southern shore of Lake Mezzola and in Pian di Spagna wetland was outlined by visual interpretation of high-resolution (<2 m pixel side) spaceborne images through the use of geographic information software (QuantumGIS v3.4.3). Satellite images used were coming from Google Earth (acquisition date: 10 April 2016) and WorldView-2 satellite (acquisition date: 01 July 2017).

## 4. Methods

Sentinel-2 data were downloaded as Top-Of-Atmosphere (TOA) reflectance data (Level-1C products). TOA data were corrected for atmospheric effect using two different algorithms, depending on the target product. For water quality retrieval, water-leaving radiance reflectance (ρw) was derived using ACOLITE, specifically developed for coastal and turbid environments and adapted to Sentinel-2 data (Vanhellemont and Ruddick, 2016), selecting the per-pixel shortwave infrared-based aerosol correction. For aquatic vegetation mapping, surface reflectance (ρ0) was derived using SEN2COR (Louis et al., 2016), including embedded topographic correction (SRTM DEM) and adjacency effect compensation (0.5 km radius).

Starting from the corrected Sentinel-2 reflectance dataset, three different mapping products were derived: i) a time series of water turbidity for Lake Mezzola; ii) benthic substrate and submerged macrophytes cover maps for the southern sector of Lake Mezzola; and iii) seasonal dynamics maps for reed beds in Pian di Spagna wetland.

### 4.1 Water turbidity maps

A time series of water turbidity maps at the surface layers for Lake Mezzola (2016-2018), expressed in formazin nephelometric units (FNU) at 20 m resolution, was produced from the ACOLITE-derived ρw, applying the single-band turbidity algorithm of Dogliotti et al. (2015), expressed in equation 1.

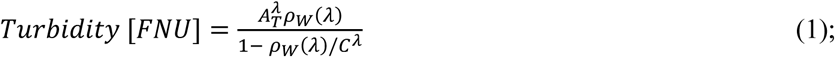

where ρw refers to Sentinel-2 spectral band 4 (red) or band 8 (near infrared), and AT and C were two wavelength-dependent calibration coefficients taken from Nechad et al. (2009) and recalibrated to Sentinel-2 spectral ranges. The algorithm is reliable over a wide range of turbidity values (1-1500 FNU), avoiding saturation issues by adopting a band-switching scheme between red and near infrared ranges (Dogliotti et al., 2015).

### 4.2 Benthic substrate maps

Maps of benthic substrate and submerged macrophytes cover in the southern sector of Lake Mezzola were derived at 10 m resolution from Sentinel-2 surface reflectance dataset, at key dates of different seasons (early August in 2017 and 2018, and late September in 2016 and 2018), using the BOMBER tool (Giardino et al., 2012). The tool implements a spectral inversion procedure of bio-optical models for both optically-deep and optically-shallow waters, based on Lee et al. (1999). BOMBER was run for estimating bottom types and cover fractions of bottom coverage, whilst setting water column properties as constant (e.g. Fritz et al., 2019). In fact, for depths up to 10 m, as in the case of the shallow bank south of Lake Mezzola, the model sensitivity to variations of water column properties is low and therefore bottom type estimation is less dependent on possible errors due to water parameters setting (Giardino et al., 2016).

Optically-shallow water areas were detected when optimization error on spectral inversion for the optically-deep model run surpassed 10% (Giardino et al., 2016 and reference herein). The bottom reflectance for the optically-shallow model was then parametrised by using the reflectance values of sand and submerged vegetation of a mix of macrophyte species (Bresciani et al., 2012), similar to those found in Lake Mezzola. The specific water absorption and backscattering coefficients were the same used by Ghirardi et al. (2019) for Lake Iseo. The concentrations of water constituents were set differently for each processed Sentinel-2 date and parameter, namely: i) suspended particulate matter values from ACOLITE turbidity outputs, by assuming a unity conversion factor (Jafar-Sidik et al., 2017); ii) coloured dissolved organic matter absorption values at 440 nm assumed fixed at 0.1 m^−1^; and iii) chlorophyll-a concentration varying according to the month, i.e. 5 mg m^−3^ for early August and 2 mg m^−3^ for late September.

### 4.3 Reed seasonal dynamics maps

Starting from the spectral bands of the Sentinel-2 surface reflectance, the three-year time series of Water Adjusted Vegetation Index (WAVI; Villa et al., 2014a) was calculated, according to equation 2. WAVI is a spectral index developed as a proxy of density and biomass of aquatic vegetation, particularly effective for emergent vegetation, i.e. helophytes (Villa et al., 2014a; 2014b).

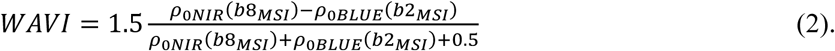

WAVI layers were produced, at 10 m spatial resolution, only for area covered by reeds in southern Lake Mezzola and Pian di Spagna, making use of the reed beds map described in Section 3.2. Areas covered by dense clouds, identified for each Sentinel-2 date as the pixels labelled with a cloud confidence level higher than 20% (i.e. SEN2COR output layer “cloud_confidence” > 20) were masked out from the WAVI time series (mask value = −1). Because S-2B data were available only from July 2017, a time series consistent across the three years was built retaining only S-2A scenes. The final WAVI series covering the three years (2016-2018) at nominal 10-day temporal resolution (revisit time of S-2A) were prepared by following Villa et al. (2018), i.e. filling missing dates with void layers (value = −1).

Seasonal dynamics metrics of reed beds in Pian di Spagna wetland were then derived running TIMESAT software (Jönsson and Eklundh, 2002; 2004) with 10-day WAVI series as input. TIMESAT was set for running with no spike filtering, Asymmetric Gaussian curves as fitting method, and two iterations for the envelope fitting (Gao et al., 2008; Villa et al., 2018). Maps of reed seasonal dynamics (10 m resolution) were produced from TIMESAT outputs for: i) the start of the growing season timing, i.e. when WAVI curve reached 0.5 of the maximum amplitude during green-up (SoS, expressed as the day of the year: DOY); ii) the end of the growing season timing, i.e. when WAVI curve decreased below the 0.5 of the maximum amplitude during senescence (EoS, as the DOY); iii) the peak WAVI reached during the season, as proxy of maximum density (WAVI_max); and iv) the area under the WAVI curve, as proxy of reed seasonal productivity (WAVI_integral).

### 4.4 Assessment of landslide impacts on lake ecosystem

Turbidity values for Lake Mezzola were extracted from Sentinel-2 derived time series. In particular, average and maximum turbidity scores for lake centre were calculated for all dates falling within the aquatic vegetation growing season, i.e. April to September (DOY 100-300), in years 2016, 2017 and 2018.

Changes in submerged macrophyte cover fraction in the optically-shallow southern sector of Lake Mezzola and downstream Lake Dascio were assessed for evaluating the impacts due to increased turbidity in the landslide aftermath. Impacts in terms of pixel-wise vegetation cover change was assessed in two different moments of the growing season: at peak of growth (early August), calculating fractional cover difference between 05 August 2018 (post-event) and 05 August 2017 (pre-event), and at the end of the growing season (late September), calculating fractional cover difference between 29 September 2018 (post-event) and 29 September 2016 (pre-event). The assessment was made separately for southern Lake Mezzola and Lake Dascio, for examining possible gradients in impact along the water flow direction.

In order to explore the impacts of landslide event on ecologically differentiated clusters, reed beds of Pian di Spagna wetland were segmented into three groups, depending on their position within the ecosystem and their ecological conditions, namely: island reeds, riparian reeds, and terrestrial reeds (see Supplementary Fig. S1). Island reeds (Isl) were identified as the areas covered by reeds which are completely surrounded by water, and are the more exposed to changes in lake level water quality. Riparian reeds (Rip) were identified as the areas covered by reeds that are spatially connected to reed beds on the shoreline and lying less than 50 m inland, being directly influenced by lake level variation. Riparian reeds were further separated into two groups: those located in Lake Mezzola (Rip_M), and those located in Lake Dascio (Rip_D). Terrestrial reeds (Ter) were identified as reed covered areas that are not spatially connected to reed beds on the shoreline and lying at least 200 m inland, being therefore considered as not directly influenced by changes in water quality and sediment deposition and therefore not suffering direct impacts from landslide events. These spatial thresholds were established based on the Lake Mezzola water level variation over the period investigated (ranging from −0.2 to + 1.1 m with respect to the average level of the Mera River, 3 km downstream of the lake; see par. 3.2). The number of 10 m pixels in Sentinel-2 derived maps and area covered by each of the four groups are shown in Supplementary Fig. S1.

Seasonal dynamics metrics (SoS, EoS, WAVI_max, WAVI_integral) for the four reed groups were extracted from satellite based maps for each of the three years (2016, 2017, 2018); from the extracted metrics data, the differences in sample distribution (*p*-value and effect size) among years (2016, 2017, 2018) separately for each reed group (Isl, Rip_M, Rip_D, Ter) were calculated.

### 4.5 Statistical analysis

Statistical analysis of differences within groups was performed using R v.3.6.1, with packages ggplot2 3.2.1, ggpubr 0.2.2, FSA 0.8.25, rcompanion 2.3.0 (Mangiafico, 2016; Wickham, 2016; Ogle et al., 2019; R Core Team, 2019). Because of non-normality of samples, multivariate differences among years were tested using non-parametric methods, i.e. Kruskal-Wallis One Way Analysis of Variance on Ranks. Post-hoc pairwise multiple comparison were performed using Dunn’s test, and relevant *p*-value scores were calculated using adjustment with the Benjamini-Hochberg method (Dunn, 1964). Effect size of pairwise sample differences were calculated using Vargha and Delaney’s A (VDA; Vargha and Delaney, 2000). VDA scores range from 0 to 1, with extreme values indicating stochastic dominance of one group over the other, and 0.5 value indicating that two groups are stochastically equal (Vargha and Delaney, 2000). VDA values < 0.29 or > 0.71 indicate large effects.

## 5. Results

### 5.1 Water turbidity patterns

Turbidity values were extracted from Sentinel-2 based maps at the lake centre location (see Fig. 3a). As time series in Fig. 3b shows, Piz Cengalo landslides of August 2017 (DOY 235-237) triggered a massive increment in Lake Mezzola turbidity that lasted until the end of October, peaking around the second half of September (DOY 267).

**Figure 3.**
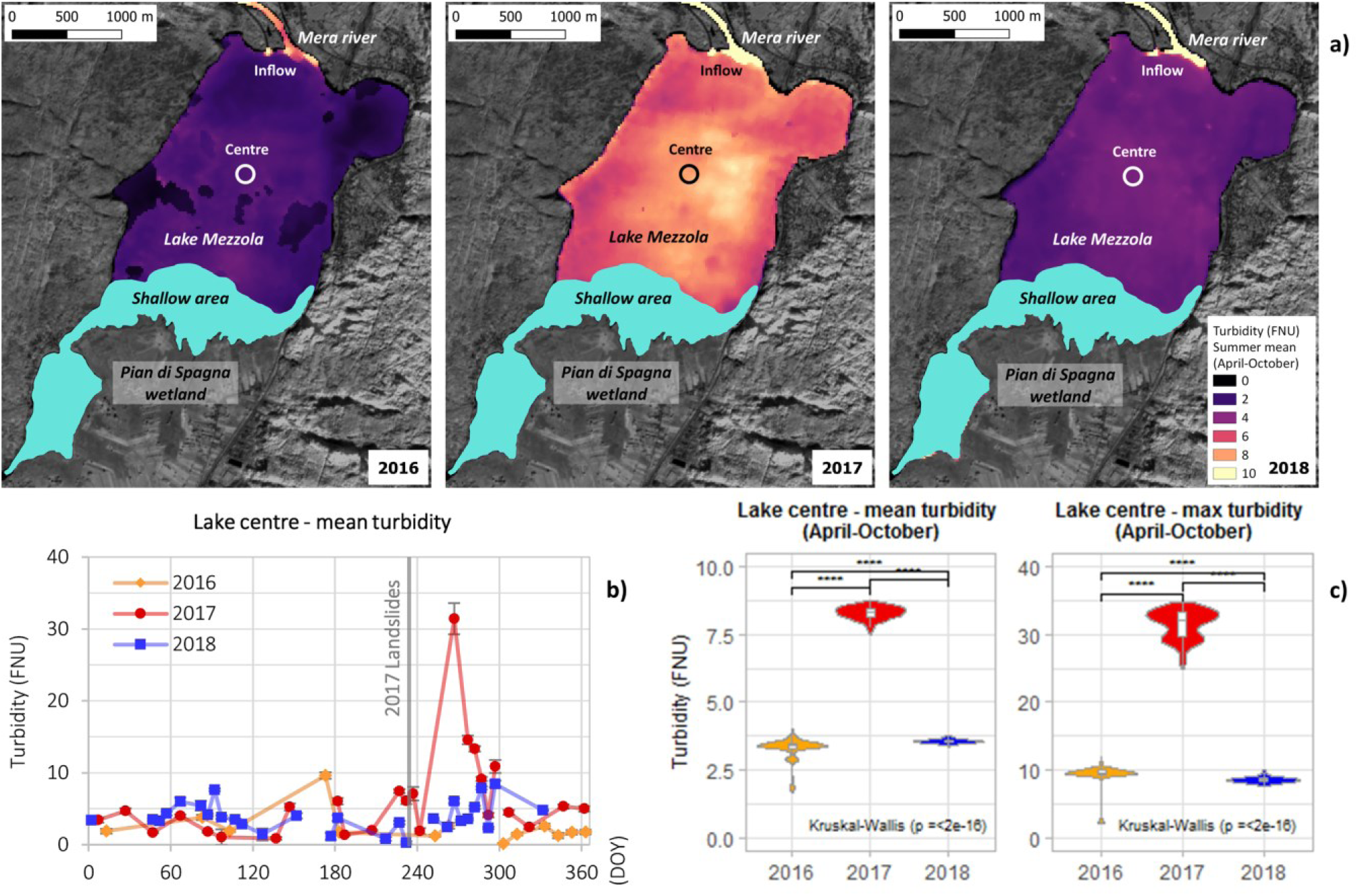
Evolution of water turbidity in Lake Mezzola derived from Sentinel-2 satellite data covering the period 2016-2018: a) maps of average water turbidity for in the macrophyte growing season (DOY 100-300) for the three years (shallow lake areas, i.e. depth < 1.5 m, are masked out); b) yearly time series of average turbidity at lake centre location extracted form satellite maps (landslide events in 2017 marked by grey vertical line); c) violin plots of average and maximum turbidity scores extracted from satellite maps during the macrophyte growing season (DOY 100-300); *p*-value scores for the differences among years (Kruskal-Wallis One Way ANOVA) and post-hoc pairwise multiple comparison (Dunn’s test) are overlaid on panel c: < 0.0001 (****); < 0.001 (***); <0.01 (*); < 0.05 (*); > 0.05 (ns).

Average turbidity for lake centre during the 2017 growing season (DOY 100 to 300), when Piz Cengalo landslides occurred, was 8.3 ± 0.2 FNU, 2.5 times the values of 2016 season, preceding the event, and 2.3 times the values of 2018 season, following it (Fig. 3c). In the same period, maximum turbidity for lake centre reached 31.5 ± 2.2 FNU in 2017, respectively 3.4 and 3.7 times the maximum of 2016 and 2018 seasons.

### 5.2 Impacts on submerged macrophytes

Submerged macrophyte cover mapped from Sentinel-2 in the southern part of Lake Mezzola showed a heavy loss in total area covered at peak of 2018 growing season, in early August, compared to the same period of the previous year, i.e. before the landslide events took place. Fig. 4a shows that change in submerged macrophyte fractional cover between 05 August 2018 and 05 August 2017 consisted in complete loss of macrophyte coverage over 28.2 ha, and partial loss of coverage over 11.7 ha, out of total 70.3 ha (partially or completely) covered by macrophytes in August 2017. Macrophyte cover at the end of growing season (29 September 2018) highlighted some signs of regain in total coverage happening in August and September 2018 (Fig. 4b): compared with pre-event conditions (29 September 2016), partial or full loss of macrophyte cover adds up to 24.6 ha, while 23.4 ha of the area increased their macrophyte fractional cover. These patterns indicate that submerged plants growth in 2018 may have been delayed compared to the previous years (hence the highest loss at early August than late September) and/or a possible shift in community composition may have happened (connected to localised partial coverage increment with respect to pre-event conditions), favouring species characterized by late vegetative peak. Indeed, during the 2018 summer, the aquatic vegetation turned out to be largely dominated by dense stands of *C. virgata*, with only a few spots of *G. densa* and *P. perfoliatus*. Moreover, the broad bare sediments were largely loosely colonized by charophyte seedlings with coverage rates below 5%.

**Figure 4.**
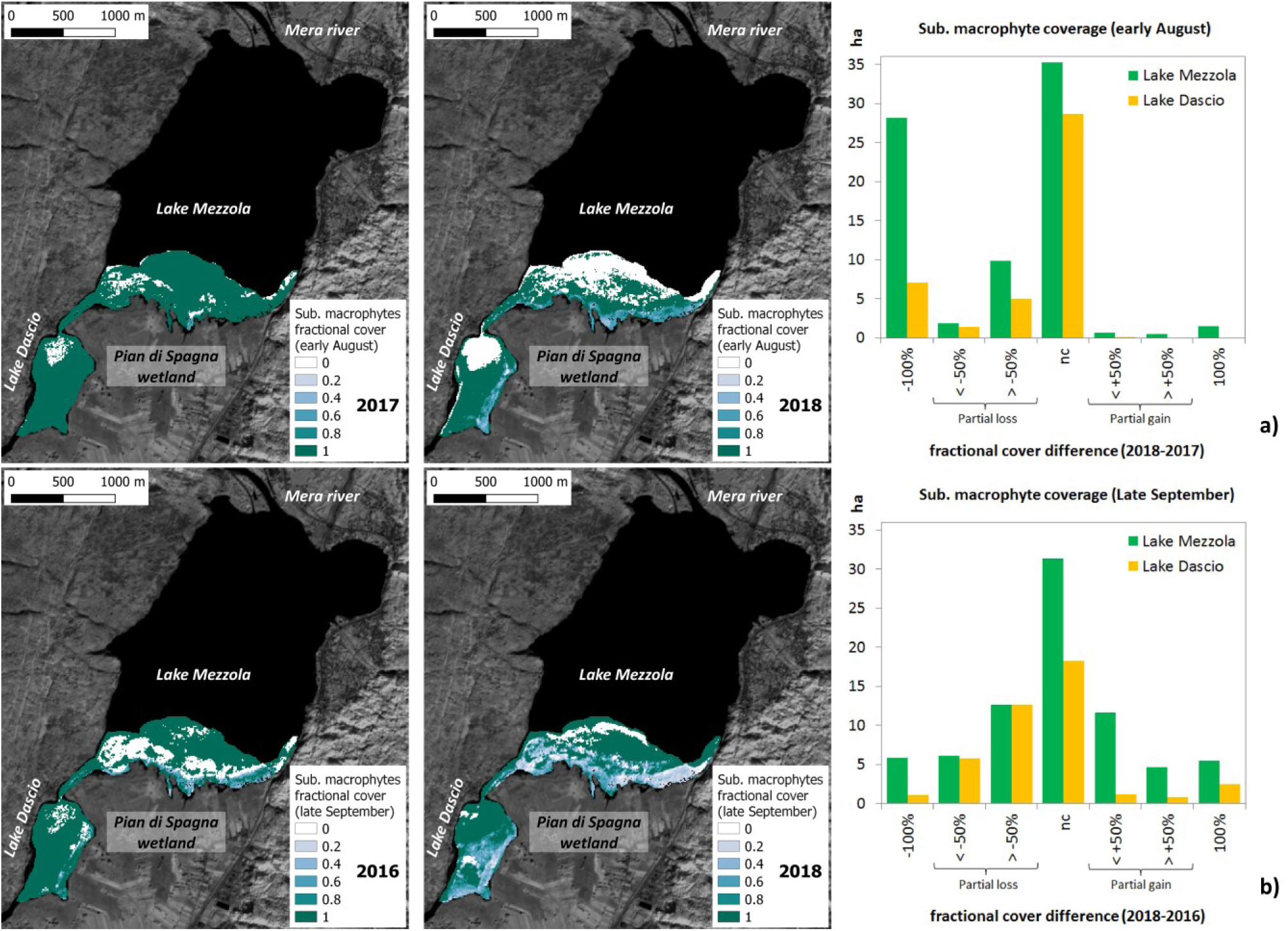
Changes in submerged macrophytes cover for southern Lake Mezzola and Lake Dascio derived from selected Sentinel-s satellite scenes: a) map of early August conditions, i.e. peak of vegetative season, in 2017 (pre-landslides) and 2018, and histograms of fractional cover difference extracted from these maps over the two lakes; b) map of late September conditions, i.e. end of growing season, in 2016 (pre-landslides) and 2018, and histograms of fractional cover difference extracted from these maps over the two lakes.

Some impacts in terms of macrophyte cover loss is visible also for Lake Dascio, although with overall magnitude far lower than what observed for upstream Lake Mezzola.

### 5.3 Impacts on riparian reed beds

Satellite-based maps of seasonal dynamics of reed beds in Pian di Spagna wetland revealed evidence of strong stress conditions for riparian reeds, in terms of green-up (SoS, Fig. 5a), and green-down (EoS, Fig. 5b) timing, as well as seasonal productivity (WAVI_integral, Fig. 5d).

**Figure 5.**
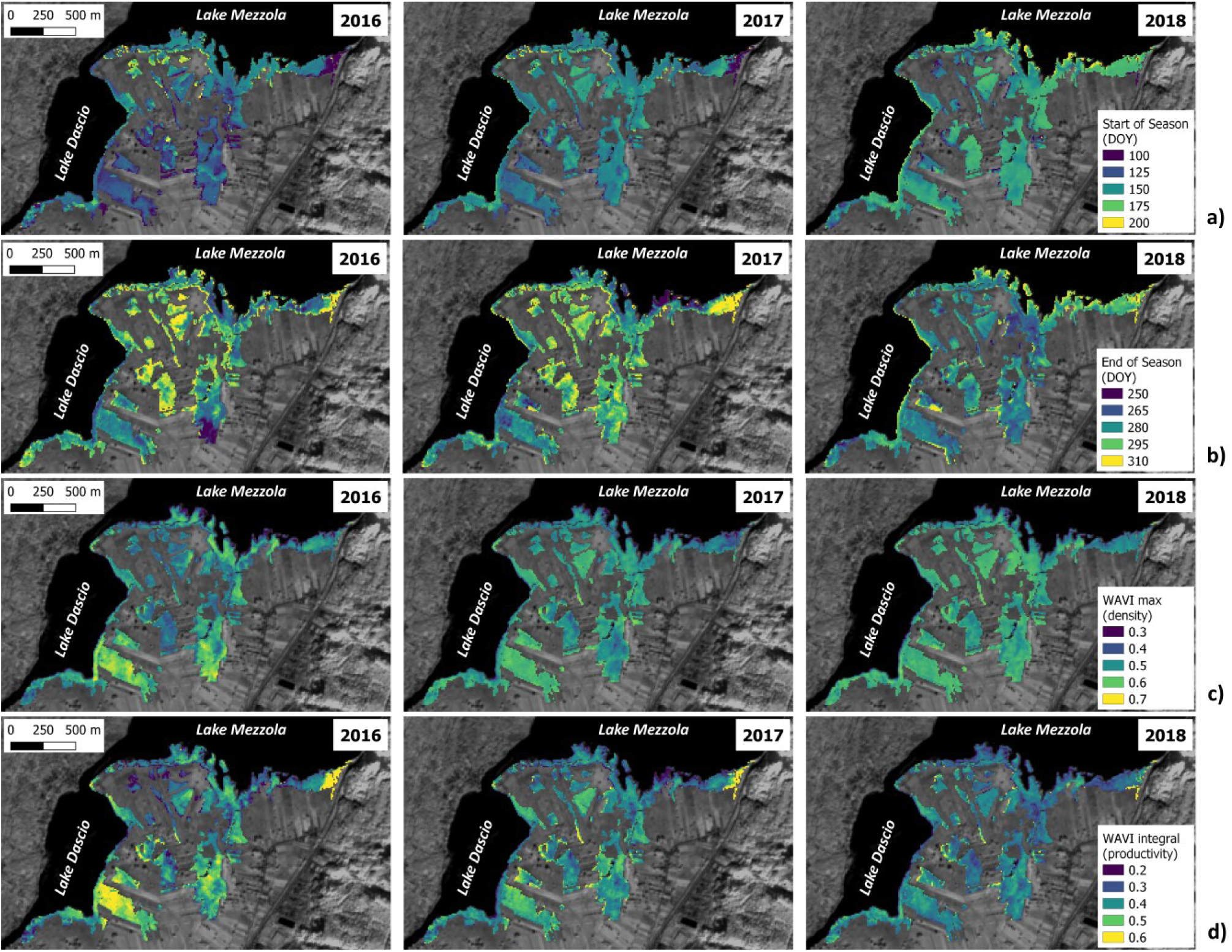
Maps of seasonal dynamics of reed beds in Pian di Spagna wetland derived from Sentinel-2 satellite time series for the three years from 2016 to 2018: a) start of season (SoS); b) end of season (EoS); c) a spectral proxy of peak canopy density (WAVI_max); d) a spectral proxy of seasonal productivity (WAVI_integral).

SoS of reed beds in Lake Mezzola came later in 2018 than in previous seasons by 30-33 days for island reeds (Fig. 6a; *p*<0.0001; VDA>0.916) and by 20-22 days for riparian reeds (Fig. 6a; *p*<0.0001; VDA>0.841). Right panel of Fig. 5a, relative to 2018 season, clearly show that such delay is particularly evident, i.e. up to 50 days later than in 2017, for reeds lying within the first 10-50 m from the waterfront and located in central part of Lake Mezzola southern shore. SoS for terrestrial reeds (rightmost panel of Fig. 6a) was progressively delayed from 2016 (DOY 128 ± 18) to 2017 (DOY 147 ± 11), and 2018 (DOY 155 ± 16).

**Figure 6.**
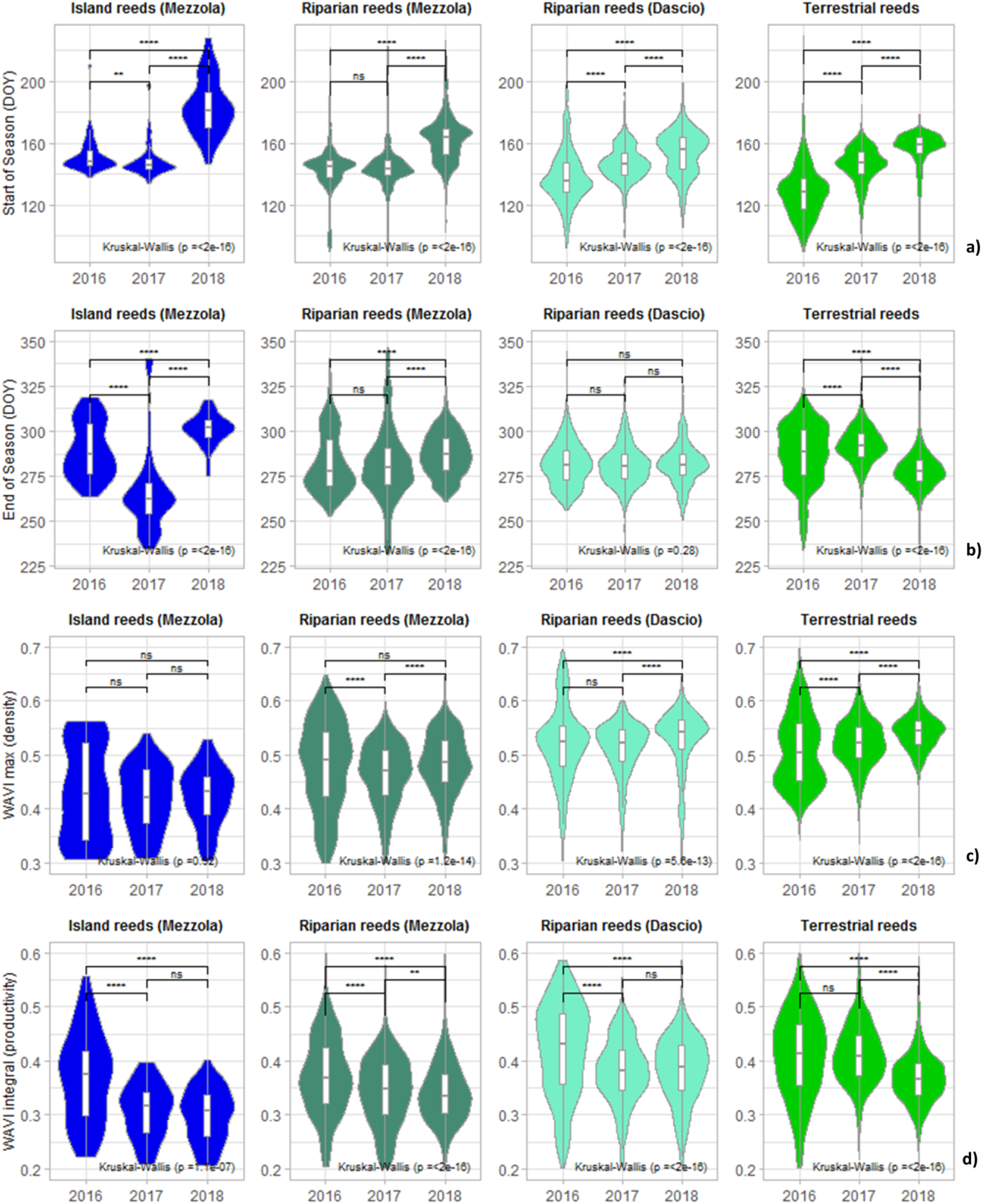
Violin plots (with encompassed box plots) of reed beds seasonal dynamics across the three years (2016-2018) for each separate reed group, i.e. Island reeds, Riparian reeds (in Lake Mezzola and Dascio), and Terrestrial reeds: a) start of season (SoS); b) end of season (EoS); c) a spectral proxy of peak canopy density (WAVI_max); d) a spectral proxy of seasonal productivity (WAVI_integral); *p*-value scores for the differences among years (Kruskal-Wallis One Way ANOVA) and post-hoc pairwise multiple comparison (Dunn’s test) are overlaid on panel c: < 0.0001 (****); < 0.001 (***); <0.01 (*); < 0.05 (*); > 0.05 (ns).

EoS for island reeds in 2017, after landslide happened, came earlier by 24 and 35 days, respectively compared to 2016 and 2018 (Fig. 6b; *p*<0.0001; VDA<0.156), while riparian reeds in both lakes Mezzola and Dascio on average did not show notable differences in green-down timing across the three years (Fig. 6b; 0.371<VDA<0.517). The spatial patterns of 2017 EoS depicted in central panel of Fig. 5b and violin plots of Fig. 6b (2^nd^ panel from the left), tough, show that this average condition for Lake Mezzola reeds is a composite effect of early senescence (around DOY 250) for riparian reeds located in central part and late senescence (around DOY 320) for those growing in easternmost part of southern lake shore. An advance of 22 days on average for EoS of terrestrial reeds was observed in 2018, compared to 2017 season (Fig. 6b; *p*<0.0001; VDA=0.156).

WAVI_integral of island reeds for both 2017 and 2018 seasons was lower than in 2016, respectively by 21% and 19% (leftmost panel of Fig. 6d; *p*<0.0001; VDA<0.272). Riparian reeds in both lakes Mezzola and Dascio did not show such highly sensitive differences in this proxy of seasonal productivity across the three years, although some minor effect in terms of lowered WAVI_integral is visible for Mezzola when 2018 (post-event) and 2016 (pre-event) conditions are compared (2^nd^ panel from the left of Fig. 6d; VDA=0.643). Terrestrial reeds experienced a loss in WAVI_integral for 2018 with respect to 2017 by 11% (rightmost panel of Fig. 6d; VDA=0.731). Out of the three seasons we considered, 2018 was generally the less productive for helophytes of Pian di Spagna wetland (Fig. 5d and 6d).

No major difference was instead observed across the years for WAVI_max of any considered reed group (Fig. 6c; 0.337<VDA<0.564) even if some local patterns are visible in Fig. 5c. This indicates that maximum canopy density reached during the growing season (surrogated by WAVI) was somewhat stable in 2016-2018 in spite of changing meteorological drivers (see Supplementary Fig. S3) as well as landslide aftermaths.

The complete picture of results from statistical test of differences in seasonal dynamics metrics among years carried out for each reed group as well as their effect size (VDA) is given in Supplementary Table S1.

## 6 Discussion

Satellite multi-temporal data provided a useful source of information for assessing ecosystem state in a perialpine lake, in terms of water quality and macrophytes, both on cover and health conditions. Monitoring capabilities demonstrated by the presented approach were amplified by the specific characteristics of Sentinel-2, in terms of data revisit (5-10 days), spatial resolution (up to 10 m) and spectral content extending from visible to shortwave infrared range, that meet the requirements for effective, operational ecosystem monitoring (Kutser et al., 2006; Song et al., 2009).

Operational mid-resolution satellite data have recently been used for environmental applications on aquatic systems focusing, for example on: riparian vegetation classification (Stratoulias et al., 2015), mapping water quality and submerged macrophyte cover (Dörnhöfer et al., 2016; Fritz et al., 2019), estimating aquatic vegetation biomass (Gao et al., 2017), assessing the spatio-temporal evolution of primary producers (Pinardi et al., 2018; Ghirardi et al., 2019), analysing the seasonal dynamics of wetland plant communities (Villa et al., 2018), and mapping cyanobacteria blooms (Sòria-Perpinyà et al., 2020).

Our work further shows the capabilities of mid-resolution satellite data and enlarges the scope with respect to previous studies, by providing a comprehensive application that joins water quality (turbidity) and aquatic to wetland vegetation (submerged macrophytes and helophytes) monitoring. In particular, the actual contribution of satellite observations to freshwater ecosystem monitoring was demonstrated for assessing the response to an extreme natural event on a real case. Sentinel-2 derived maps allowed us to highlight the turbidity patterns in Lake Mezzola in space and time, the evolution of submerged macrophyte cover in different years and the quantitative analysis of seasonal dynamics of helophytes communities living on the lake shores. The approach we proposed could be easily implemented for continuous monitoring of Lake Mezzola and surrounding areas during the current and next years, so to verify the evolution of aquatic vegetation conditions and ecosystem equilibrium towards recovery or further degradation. Moreover, further applications could be potentially extended to similar ecosystems - with water bodies larger than 2 ha and hosting macrophyte communities with individual patches larger than 100 m^2^-susceptible to periodically extreme sediment loads, which across the alpine and peri-alpine region could be due to upstream dam regulations or landslides. The August 2017 landslides triggered an increase in turbidity that amounted to twice the average and three times the maximum values scored in the years preceding and following the events. Satellite-based maps revealed that river inflow in northern part of the lake and the lake central part are hotspots of turbidity, highlighting surface circulation patterns. However, during 2017, high values of turbidity were scored also for southern portion of Lake Mezzola, where a shallow bank (depth < 7 m) hosts abundant macrophyte communities, and the helophyte beds of Pian di Spagna wetland reach the open waters.

Increased turbidity is a direct consequence of the massive load of detritus brought by the Mera River after the Piz Cengalo landslides. Material brought by Mera River, almost completely inorganic (mainly made by a mixture of silica and alumina, see Pettine et al., 2000), sedimented in Lake Mezzola and was the cause of serious impacts on the lake ecosystem. The impacts shown by analysing the time series of Sentinel-2 derived products were particularly intense on submerged macrophytes and riparian helophytes in southern part of the lake, with cascade effects on the whole trophic chain and animal communities relying on them as primary habitat.

Light availability is one of the key factors affecting submerged macrophyte establishment and productivity (e.g. Barko et al., 1986, Lacoul and Freedman, 2006), with implications for growth rates and plant morphology (e.g. Riis et al., 2012). High levels of turbidity and sediment deposition are therefore connected with negative impacts on benthic vegetation, because they reduce light availability for photosynthesis and can physically hinder germination of seeds or sprouting of new buds at the beginning of the season (Wood and Armitage, 1997; Spencer and Ksander, 2002).

Our results documented a partial loss of total cover of submerged macrophytes (including charophyte beds) in the season following the landslides, accompanied by signs of late growth, with the coverage peak moving towards late September. Moreover, the analysis of spatial clustering of benthic vegetation dynamics shown in Fig. 4 suggests the possibility of shifts in community dominance in favour of species less sensitive to the effects of abundant deposition of inorganic, nutrient-poor sediments. Our 2018 surveys highlighted a clear predominance of *C. virgata* over *G. densa* and *P. perfoliatus*. An annual or perennial species depending on the growing depth, *C. virgata* can tolerate competition from other water plants (Guiry, 2019). Indeed, in newly colonized or recently perturbed habitats charophytes may exhibit a strong pioneer behaviour and be highly competitive towards vascular species (Bonis and Grillas, 2002; Brochet et al., 2010), despite of their disadvantage in non-oligotrophic environments (Blindow, 1992). Our observations seem to go in this direction, and we could explain the widespread presence of *C. virgata* in the southern banks of Lake Mezzola referring to this pioneering habit. We therefore hypothesize that the submerged macrophyte cover mapped after the events in 2018 could represent an initial stage of recolonization after a strong perturbation. This process may have been strengthened by the physical and chemical features of particulate material brought in the lake as a consequence of the landslides. The characteristics of this detrital material, rich in silica and alumina, could support the widespread presence of oligotrophic species (i.e. charophytes and isoetids), otherwise rare in areas tending to mesotrophic conditions (see Blindow, 1992; Sand-Jensen et al. 2008), such as Lake Mezzola is. Such an effect of landslide deposits on submerged macrophyte community composition could bring along interesting implications from the point of view of environmental management (e.g. of mountain reservoirs and lakes), but requires further verification.

While temperature is considered the main factor controlling phenology of common reed (Irmak et al., 2013; Petus et al., 2013; Anda et al., 2017), our results show that exceptional sediment load and deposition (e.g. due to upstream landslides) can overwrite climatic factors in determining reed seasonal dynamics change. Specifically, riparian reeds located south of Lake Mezzola suffered from an anticipated senescence in 2017 (late September, on average), initiated in the weeks after Piz Cengalo landslides, around 20 days before that of 2016, as well as from a delayed green-up by more than 20 days in the following growing season, i.e. with average SoS for island reeds in early July 2018.

Although 2018 was characterized by unusually cold early spring in North Italy, with growing degree-days (GDD) cumulated up to mid-April lower than 1995-2014 average by 26°C d (−22% in relative terms, see Supplementary Fig. S3), the late development of riparian reed beds of Lake Mezzola in 2018 (+33.2 and +21.9 days on average in SoS compared to 2017 situation, for island and riparian stands respectively) was not attributable only to meteorological anomaly. In fact, terrestrial reeds of Pian di Spagna wetland, which we supposed not to be impacted by deposition of sediments in landslide aftermaths, due to their distance from the shoreline (> 200 m), showed themselves some signs of delayed growth in 2018 (+9.2 days in SoS on average, compared to 2017 conditions), but with far lower magnitude. Consistently with this, Anda et al. (2017) found low temperatures in spring to delay reed growth by one week in Kis Balaton wetland, in Hungary (46°37’ N, 17°08’ E). In addition to temperature in early spring, the extremely dry spring of 2018, with −21% total cumulated rainfall from January to June, compared to 1995-2014 average (Supplementary Table S2), might have contributed to slightly delayed SoS for terrestrial reeds, more sensitive to water stress (Haslam, 1970).

The impact of landslide-triggered sediment deposition was evident already in autumn 2017 on island reed communities, which suffered from senescence anticipated by three weeks with respect to pre-event conditions (2016). In 2018, the situation went back to normality (average EoS at DOY 301), also favoured by hot weather lasting longer than usual (mean temperature in October higher by 2.7°C compared to 1995-2014 average, see Supplementary Fig. S3). On the other hand, the prolonged drought conditions of 2018 season, with −22% total cumulated rainfall from January to September compared to 1995-2014 average (Supplementary Table S2), most probably amplified the water stress of terrestrial reed communities (Haslam, 1970), driving to an advanced green-down, about 10 days earlier than 2016 and 2017.

The anticipated senescence of 2017 and later green-up of 2018 resulted in two consecutive years of reduced seasonal productivity (around −20% in terms of WAVI_integral scores) for riparian reed beds of Lake Mezzola, with possible serious consequences in terms of reserves stored in rhizomes and capabilities to resist further catastrophic events (Čížková et al., 2001).

Satellite-based maps are inevitably affected by a certain degree of uncertainty, due to the inherent complexity of targets, i.e. biological and ecological systems. Water turbidity products derived with the method implemented as in Section 4.1 have been assessed by Dogliotti et al. (2015) using *in situ* data covering different sites and a wide range of turbidity (1-1000 FNU), scoring a mean relative error around 13.7%. Accuracy of benthic substrate cover derived with the method described in Section 4.2 has been assessed by Ghirardi et al. (2019) over perialpine Lake Iseo against *in situ* reference data, resulting in per-class accuracies (F-score) of 96.6% and 87.5% for submerged macrophytes and bare sediment cover classes, respectively. Possible bias in key metrics of macrophyte seasonal dynamics derived from 10-day revisit Sentinel-2 time series as delineated in Section 4.3 have been estimated with respect to the best temporal resolution available for mid-resolution operational satellites by Villa et al. (2018) over emergent and floating plants of Mantua lakes system, as under 2 days or 3 days for SoS and EoS, respectively.

The aforementioned uncertainty levels expected for the three ecosystem parameters at the basis of our analysis do not substantially bias the primary findings described above, as they are generally around one order of magnitude lower than the main effects of landslide impacts on Lake Mezzola ecosystem: i.e. water turbidity increased by 2.3 to 3.7 times due to the landslide, and 20-30 day differences in SoS and EoS measured for riparian reeds after the landslide aftermaths.

The impacts on riparian reeds are most visible in reed beds completely surrounded by water, thus more sensitive to changes in water quality, as shown in Fig. 5 maps, i.e. representing SoS for 2018 (Fig. 5a, right panel) and EoS for 2017 (Fig. 5b, central panel). Traces of dead, broken stems observed *in situ* in late September 2018 (Supplementary Fig. S2) witnessed a retreat of the reed front with respect to previous maximum coverage, as well as reed canopy thinning of some areas in central southern shore of Lake Mezzola, particularly impacted according to Fig. 5 maps, thus corroborating satellite-based findings.

The decoupling between seasonal productivity proxy WAVI_integral, clearly showing the consequences of landslides impacts on riparian reeds in both 2017 and 2018, and density-sensitive WAVI_max, not showing sensitive differences among the three years for all reed types, could be explained by the growth of a turf populated mainly by isoetids and other amphibious species - *L. uniflora, E. acicularis, L. aquatic* - below the reeds in 2018 summer (documented during the *in situ* survey of 26 September 2018, see Supplementary Fig. S2). The expansion of this understory was probably promoted by landslide impacts in a dual way, that is: i) deposition of materials and increase in turbidity have weakened the riparian reed, as traces of broken reed straws and low density reed patches were found during the *in situ* survey would support; and ii) the inorganic sediments brought by the Mera river into the lake favoured the development of species adapted to organic-poor substrates, as already observed for submerged macrophytes. Expansion of the understory and thinning of riparian reeds probably balanced one each other in terms of integrated spectral response at canopy scale, resulting in 2018 peak WAVI scores similar to those of previous years, while degradation of riparian reed communities after 2017 summer landslides came out evidently from other seasonal dynamics metrics.

A possible explanation of the above described impacts of landslide aftermath on riparian reed belt of Lake Mezzola can be found not only in the physical action of increased sediment loadings hampering rhizomes oxygenation, but also in the quality of such new sediments, rich in silica and alumina. In fact, even if low to moderate enrichment with silica can provide benefits for reed growth (Máthé et al., 2012; Schaller et al., 2012a), high concentrations can compromise growth, possibly by inhibiting uptake of beneficial metals, i.e. iron (Máthé et al., 2012; Schaller et al., 2012b). Literature on effects of alumina on reed plants are instead not decisive, but tend to highlight some positive effects on phosphorous sequestration resulting into competitive advantage towards other wetland species (Meyerson et al., 2002; Batty and Younger, 2004).

## Conclusions

Monitoring the evolution of key lake ecosystem variables and their response to external events using remote sensing is feasible, provided that satellite data with adequate spatial and temporal resolutions are available and an approach is implemented based on products mapping different environmental parameters in a comprehensive way.

Our work demonstrated the contribution of Sentinel-2 derived multi-temporal maps in assessing the impacts of a landslide event on the ecosystem of Lake Mezzola (Northern Italy). We found a connection between the landslide aftermath and lake ecosystem dynamics under different aspects, that are: i) water turbidity patterns and their temporal evolution; ii) loss of biomass and possible shift in species compositions for submerged macrophyte communities; iii) shortened growing season and reduced productivity for riparian reed beds on the southern lake shore, due to early senescence in 2017 and delayed start of growth in 2018.

The results have shown that, although 2018 season was anomalously dry and hot compared to the previous ones, the highlighted impact on riparian reeds is not attributable to meteorological drivers, as the terrestrial reed beds in the area did not show the same delay in green-up and loss of productivity.

The utilization of Sentinel-2 satellite data in the framework of the presented approach would make possible for environmental bodies and public authorities to carry on monitoring the state of Lake Mezzola ecosystem equilibrium, in order to assess the evolution trend whether it goes towards recovery of further degradation for both submerged macrophyte communities and helophytes, and also that of similar ecosystems subject to external pressures with periods of high sediment loads.

## Author contributions

**Paolo Villa**: Conceptualization, Methodology, Analysis, Investigation, Writing of original draft. **Mariano Bresciani**: Conceptualization, Methodology, Investigation. **Rossano Bolpagni**: Analysis, Investigation. **Federica Braga**: Methodology. **Dario Bellingeri**: Investigation. **Claudia Giardino**: Supervision. All authors reviewed and edited the final manuscript.

## Acknowledgements

The authors thank Jasmine S. Zanenga for her help in Sentinel-2 pre-processing and GIS assisted delineation of reed beds of Pian di Spagna wetland. The authors are grateful to Chiara Agostinelli and Elisa Carena (ARPA Lombardia) for enabling the *in situ* survey of Lake Mezzola southern shore of 26 September 2018. Hydro-meteorological data covering the study area were provided by ARPA Lombardia (https://www.arpalombardia.it/siti/arpalombardia/meteo/richiesta-dati-misurati/). Sentinel-2 data were provided by EU programme Copernicus (https://scihub.copernicus.eu/). This work was supported by the EU Horizon 2020 programme through the EOMORES project [grant number 730066].

