## Supplemental materials for "Impact of upstream landslide on perialpine lake ecosystem: an assessment using multi-temporal satellite data"

### Supplementary Material

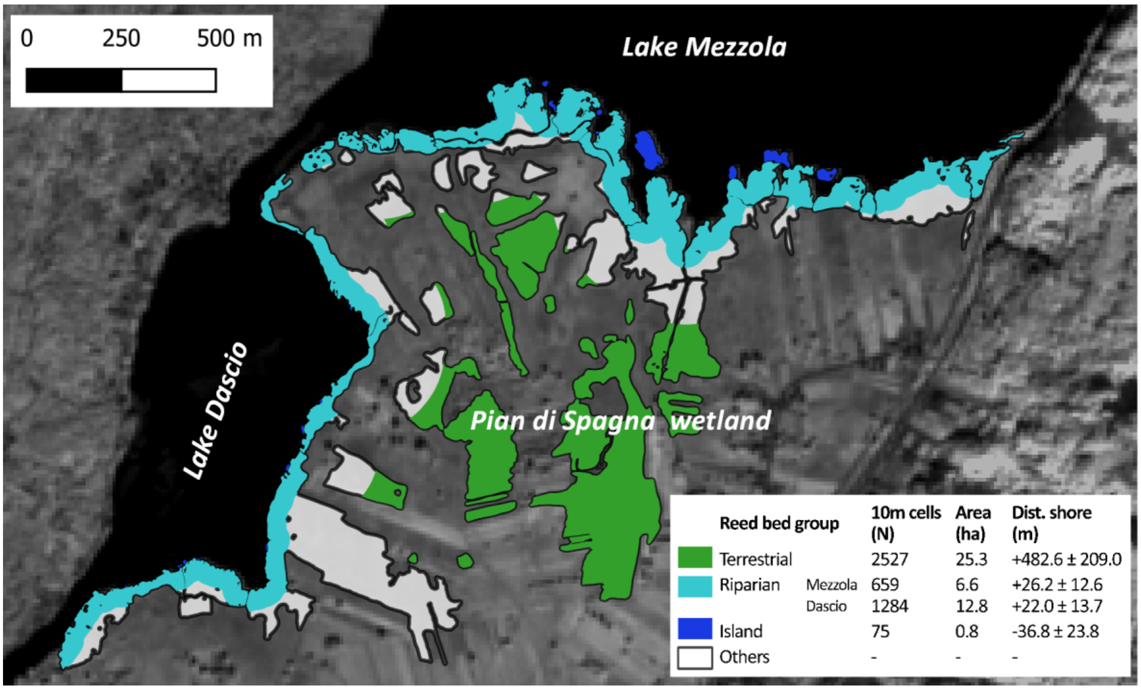


**Figure S1.** Reed beds map highlighting the different groups considered in the analysis of Piz Cengalo landslide aftermath impacts, and main features of each group, in terms of area covered, corresponding number of 10-m pixels in satellite-derived seasonal dynamics maps, and distance from lake shoreline (average ± standard deviation).

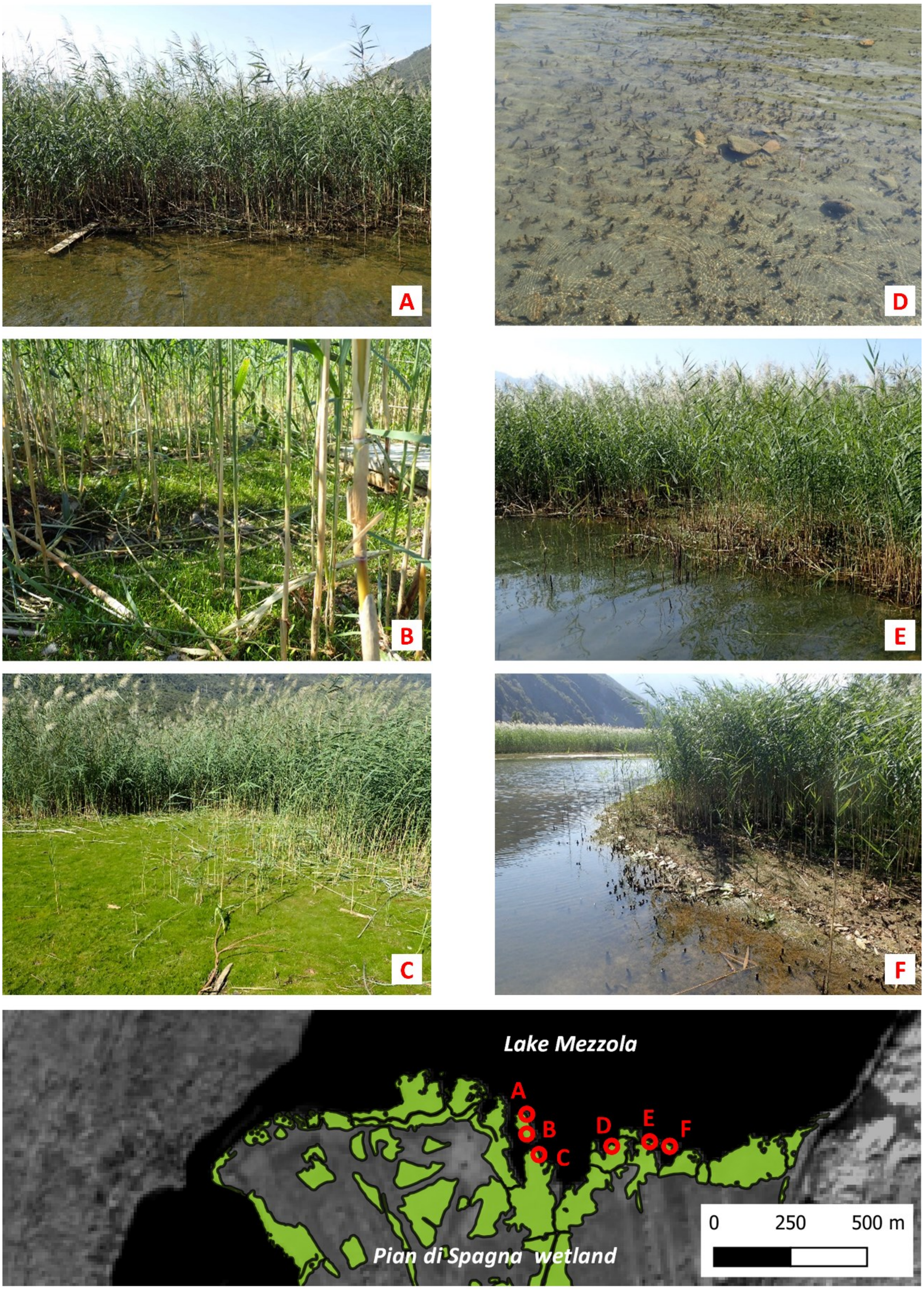


**Figure S2.** Georeferenced photos (position shown on lower panel) taken along the southern shore of Lake Mezzola during the *in situ* survey showing riparian reed communities conditions on 26 September 2018: signs of retreat of reed front with sparse, short plants (A and E), development of dense understory of short grasses species (B and C), and remnants of broken stems left from previous seasons (D and F).

**Table S1**. Statistical test of differences in seasonal dynamics metrics among years for each reed group.

**Start of Season (SoS)**

| Comparison | Z [b] | *p*-value [b] | VDA [c] |
|---|---|---|---|
| Island reeds (Mezzola) | | | |
| ANOVA [a]: $\chi^2$ = 110.9 (n=75, df=2, *p*-value<2.2e-16) | | | |
| 2016-2017 | 2.245 | 2.48E-02 | 0.652 |
| 2016-2018 | -7.789 | 1.02E-14 | 0.084 |
| 2017-2018 | -10.033 | 3.26E-23 | 0.068 |
| Riparian reeds (Mezzola) | | | |
| ANOVA [a]: $\chi^2$ = 1277.8 (n=1284, df=2, *p*-value<2.2e-16) | | | |
| 2016-2017 | 2.616 | 8.91E-03 | 0.534 |
| 2016-2018 | -29.567 | 5.99E-192 | 0.159 |
| 2017-2018 | -32.182 | 9.32E-227 | 0.137 |
| Riparian reeds (Dascio) | | | |
| ANOVA [a]: $\chi^2$ = 360.2 (n=659, df=2, *p*-value<2.2e-16) | | | |
| 2016-2017 | -10.626 | 3.40E-26 | 0.300 |
| 2016-2018 | -18.931 | 1.90E-79 | 0.229 |
| 2017-2018 | -8.305 | 9.94E-17 | 0.337 |
| Terrestrial reeds | | | |
| ANOVA [a]: $\chi^2$ = 3206.3 (n=2527, df=2, *p*-value<2.2e-16) | | | |
| 2016-2017 | -31.054 | 1.50E-211 | 0.183 |
| 2016-2018 | -56.533 | 0.00E+00 | 0.106 |
| 2017-2018 | -25.479 | 3.39E-143 | 0.228 |

**WAVI_max (canopy density)**

| Comparison | Z [b] | *p*-value [b] | VDA [c] |
|---|---|---|---|
| Island reeds (Mezzola) | | | |
| ANOVA [a]: $\chi^2$ = 1.1 (n=75, df=2, *p*-value=0.5683) | | | |
| 2016-2017 | -0.622 | 8.01E-01 | 0.472 |
| 2016-2018 | -1.058 | 8.71E-01 | 0.449 |
| 2017-2018 | -0.436 | 6.63E-01 | 0.480 |
| Riparian reeds (Mezzola) | | | |
| ANOVA [a]: $\chi^2$ = 72.6 (n=1284, df=2, *p*-value<2.2e-16) | | | |
| 2016-2017 | 6.254 | 5.98E-10 | 0.564 |
| 2016-2018 | -1.882 | 5.98E-02 | 0.485 |
| 2017-2018 | -8.136 | 1.22E-15 | 0.400 |
| Riparian reeds (Dascio) | | | |
| ANOVA [a]: $\chi^2$ = 55.6 (n=659, df=2, *p*-value=8.347e-13) | | | |
| 2016-2017 | 2.129 | 3.33E-02 | 0.524 |
| 2016-2018 | -5.126 | 4.45E-07 | 0.429 |
| 2017-2018 | -7.255 | 1.21E-12 | 0.374 |
| Terrestrial reeds | | | |
| ANOVA [a]: $\chi^2$ = 513.9 (n=2527, df=2, *p*-value<2.2e-16) | | | |
| 2016-2017 | -6.894 | 5.43E-12 | 0.427 |
| 2016-2018 | -22.149 | 3.18E-108 | 0.337 |
| 2017-2018 | -15.256 | 2.27E-52 | 0.359 |

**End of Season (EoS)**

| Comparison | Z [b] | *p*-value [b] | VDA [c] |
|---|---|---|---|
| Island reeds (Mezzola) | | | |
| ANOVA [a]: $\chi^2$ = 90.0 (n=75, df=2, *p*-value<2.2e-16) | | | |
| 2016-2017 | 6.024 | 2.55E-09 | 0.844 |
| 2016-2018 | -3.336 | 8.49E-04 | 0.284 |
| 2017-2018 | -9.360 | 2.39E-20 | 0.112 |
| Riparian reeds (Mezzola) | | | |
| ANOVA [a]: $\chi^2$ = 156.7 (n=1284, df=2, *p*-value<2.2e-16) | | | |
| 2016-2017 | 0.721 | 4.71E-01 | 0.507 |
| 2016-2018 | -10.463 | 1.91E-25 | 0.382 |
| 2017-2018 | -11.184 | 1.47E-28 | 0.371 |
| Riparian reeds (Dascio) | | | |
| ANOVA [a]: $\chi^2$ = 2.6 (n=659, df=2, *p*-value=0.277) | | | |
| 2016-2017 | 1.158 | 3.70E-01 | 0.517 |
| 2016-2018 | -0.380 | 7.04E-01 | 0.496 |
| 2017-2018 | -1.538 | 3.72E-01 | 0.474 |
| Terrestrial reeds | | | |
| ANOVA [a]: $\chi^2$ = 1567.8 (n=2527, df=2, *p*-value<2.2e-16) | | | |
| 2016-2017 | -12.907 | 4.12E-38 | 0.424 |
| 2016-2018 | 25.964 | 1.90E-148 | 0.682 |
| 2017-2018 | 38.871 | 0.00E+00 | 0.844 |

**WAVI_integral (seasonal productivity)**

| Comparison | Z [b] | *p*-value [b] | VDA [c] |
|---|---|---|---|
| Island reeds (Mezzola) | | | |
| ANOVA [a]: $\chi^2$ = 31.9 (n=75, df=2, *p*-value=1.196e-07) | | | |
| 2016-2017 | 5.051 | 1.32E-06 | 0.736 |
| 2016-2018 | 4.711 | 3.70E-06 | 0.728 |
| 2017-2018 | -0.340 | 7.34E-01 | 0.480 |
| Riparian reeds (Mezzola) | | | |
| ANOVA [a]: $\chi^2$ = 161.2 (n=1284, df=2, *p*-value<2.2e-16) | | | |
| 2016-2017 | 8.745 | 3.34E-18 | 0.598 |
| 2016-2018 | 12.344 | 1.57E-34 | 0.643 |
| 2017-2018 | 3.599 | 3.19E-04 | 0.539 |
| Riparian reeds (Dascio) | | | |
| ANOVA [a]: $\chi^2$ = 103.4 (n=659, df=2, *p*-value<2.2e-16) | | | |
| 2016-2017 | 9.299 | 4.24E-20 | 0.644 |
| 2016-2018 | 8.207 | 3.40E-16 | 0.635 |
| 2017-2018 | -1.092 | 2.75E-01 | 0.478 |
| Terrestrial reeds | | | |
| ANOVA [a]: $\chi^2$ = 803.7 (n=2527, df=2, *p*-value<2.2e-16) | | | |
| 2016-2017 | -3.277 | 1.05E-03 | 0.493 |
| 2016-2018 | 22.748 | 2.25E-114 | 0.665 |
| 2017-2018 | 26.025 | 7.74E-149 | 0.731 |

[a] among group difference tested with Krustal-Wallis ANOVA on ranks;
[b] pairwise multiple comparisons (Dunn's test, Benjamini-Hochberg adjustment);

[c] pairwise effect size calculations using Vargha and Delaney's A (VDA).

### Hydro-meteorological indicators

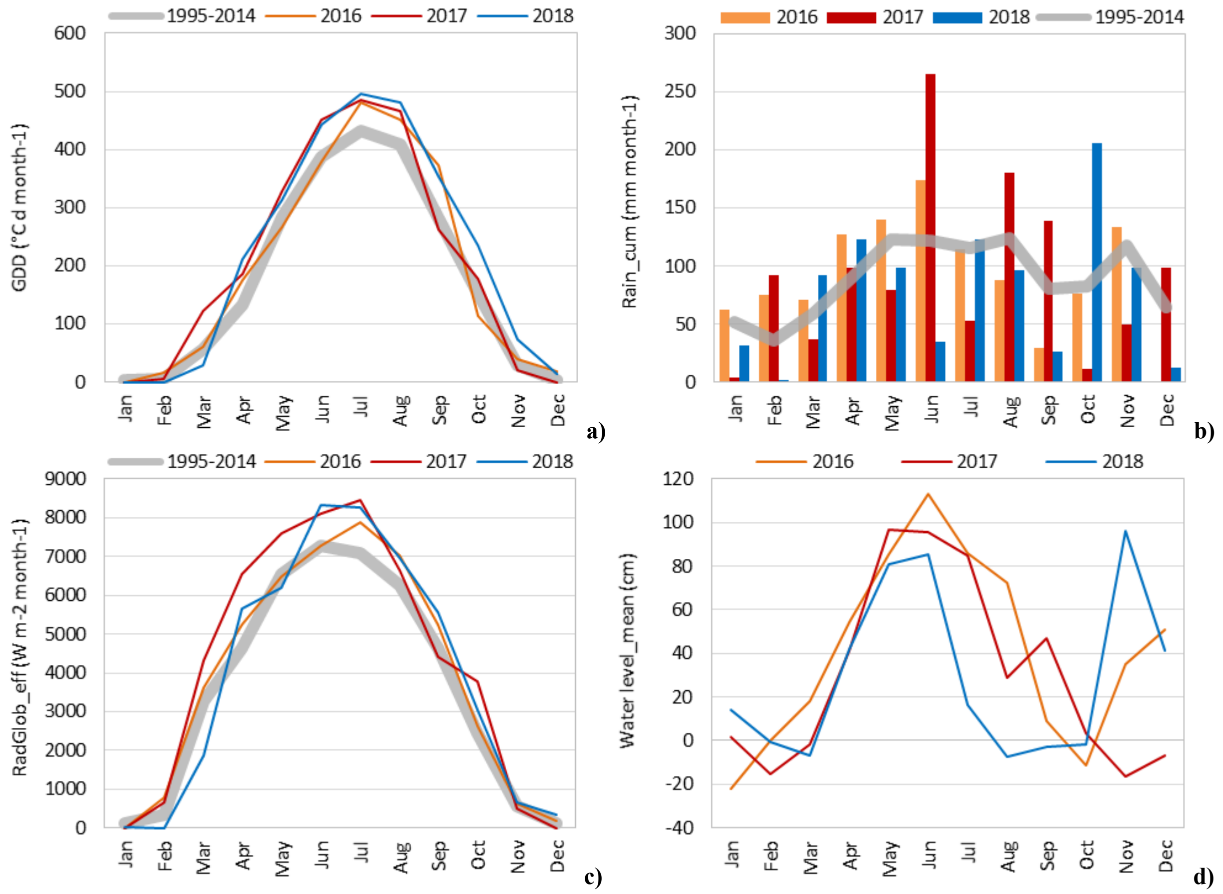


**Figure S3.** Monthly synthesis of selected hydrological and meteorological indicators recorded by Samolaco weather station, 4 km upstream of Lake Mezzola (compared to 1995-2014 average): a) monthly cumulated growing degree-days (GDD) for common reeds; b) monthly cumulated rainfall (Rain_cum); c) monthly effective cumulated global radiation (RadGlob_eff); d) average water level of the Mera River (3 km downstream of Lake Mezzola).

**Table S2.** Seasonal meteorological conditions on Lake Mezzola area for 2016-2018

| Meteorological indicator | Unit | Period | 2016 | | 2017 | | 2018 | |
|---|---|---|---|---|---|---|---|---|
| | | | value | anomaly [f] | value | anomaly [f] | value | anomaly [f] |
| Water level [a], average | cm | Mar-Apr [d]<br>Apr-Sep | 35.9<br>69.9 | na [g] | 19.4<br>65.4 | na [g] | 17.2<br>35.4 | na [g] |
| GDD [b], cumulated | °C d | Jan-Jun [e]<br>Jan-Sep | 897<br>2200 | +3%<br>+9% | 1094<br>2309 | +25%<br>+14% | 997<br>2327 | +14%<br>+15% |
| RadGlob_eff [c], cumulated | kWh $m^{-2}$ | Jan-Jun [e]<br>Jan-Sep | 562<br>1045 | +9%<br>+7% | 653<br>1123 | +27%<br>+15% | 530<br>1029 | +3%<br>+5% |
| Rainfall, cumulated | mm | Jan-Jun [e]<br>Jan-Sep | 649<br>880 | +35%<br>+10% | 575<br>947 | +20%<br>+18% | 381<br>629 | -21%<br>-22% |
| | | | Wet and mild season; optimal conditions for vegetation growth | | Wet and hot season, high light availability; optimal conditions for hydrophytes growth | | Dry and hot season; critical conditions for terrestrial vegetation growth | |

[a] water level of the Mera River, measured 3 km downstream of Lake Mezzola ouflow;
[b] growing degree-days cumulated during the period (baseline temperature of 7 °C for common reed, according to Anda et al. (2017));
[c] effective global radiation cumulated during the period (radiation computed only when daily GDD > 0 °C d);
[d] water level average at two different time scales (Bresciani et al., 2011): reed early growth (Mar-Apr: 01 March–30 April), and during the whole growing season (Apr-Sep: 01 April–30 September);
[e] key seasonal windows connected to with reed phenology: early vegetative (Jan-Jun: 01 January–30 June), and late reproductive (Jan-Sep: 01 January–30 September);
[f] anomalies are calculated as relative differences (%) with respect to average 1995-2014 conditions;

[g] not available (water level measures started in 2013).
